# Porcine Nasal Organoids as a model to study the interactions between the swine nasal microbiota and the host

**DOI:** 10.1101/2024.08.13.606910

**Authors:** Laura Bonillo-Lopez, Noelia Carmona-Vicente, Ferran Tarrés-Freixas, Karl Kochanowski, Jorge Martínez, Mònica Perez, Marina Sibila, Florencia Correa-Fiz, Virginia Aragon

## Abstract

Interactions between the nasal epithelium, commensal nasal microbiota, and respiratory pathogens play a key role in respiratory infections. Currently there is a lack of experimental models to study such interactions under defined *in vitro* conditions. Here, we developed a Porcine Nasal Organoid (PNO) system from nasal tissue of pigs as well as from cytological brushes. PNOs exhibited similar structure and cell types than the nasal mucosa, as evaluated by immunostaining. PNOs were inoculated with porcine commensal strains of *Moraxella pluranimalium, Rothia nasimurium* and the pathobiont *Glaesserella parasuis* for examining host-commensal-pathogen interactions. All strains adhered to the PNOs, although at different levels. *M. pluranimalium* and *G. parasuis* strains stimulated the production of proinflammatory cytokines, whereas *R. nasimurium* induced the production of IFNγ and diminished the proinflammatory effect of the other strains. Overall, PNOs mimic the *in vivo* nasal mucosa and can be useful to perform host-microbe interaction studies.

**In brief:** Interactions between nasal epithelium, commensal nasal microbiota, and respiratory pathogens play a key role in respiratory infections. We developed porcine nasal organoids to mimic the nasal mucosa and use as a model to study those interactions. Additionally, this development supports the reduction of the number of animals for animal experimentation.

**Highlights:**

- First generation of Porcine Nasal Organoids (PNOs) from nasal turbinates and swabs
- PNOs recapitulate features of *in vivo* tissue and maintain self-renewal capacity
- Host-microbe interactions can be studied using the PNO system
- *Rothia nasimurium* inhibits the inflammation induced by other bacteria

## INTRODUCTION

The nasal epithelium is the first barrier in direct contact with the external environment providing protection from viral, bacterial and fungal pathogens, as well as pollutants in the air^1,2^. This tissue is responsible for mucociliary clearance and, together with the nasal microbiota, acts as a physical, chemical, and immunological defence to preserve the homeostasis of the nasal cavity and limit the invasion of microorganisms^3–10^. It is also considered an integral part of the innate immune system as it produces antimicrobial agents, danger-associated molecular patterns (DAMPs) and recognize pathogens via pattern recognition receptors (PRRs)^4^. Integrity alteration of the nose epithelium has been related to inflammation with severe consequences for animal health^3,4,11^.

Traditionally, models to study basic and clinical aspects of the airway pathophysiology have been cells and explants from nose, trachea, or bronchia. These models were either hard to maintain *in vitro* (like primary cells and explants) or not physiologically relevant enough (like immortalised cell lines)^12–16^. The recent emergence of both human and animal organoids are promising systems to model the respiratory epithelium and study immunological parameters or host-pathogen interactions. To date, airway organoids established in humans^17–19^ as well as in some animal species^20–22^ have shown to represent a robust system for airway modelling, since they retain the *in vivo* architecture of the nasal epithelium forming a complex pseudostratified epithelium with most of the cell types from the original tissue (ciliated, mucus-secreting goblet, basal and club cells)^1,2^. To our knowledge, only lung organoids have been set up for the porcine respiratory tract^21^. Despite the burden of respiratory diseases for the porcine sector, there is no nasal organoid model currently available for pigs, which would enable experiments under defined *in vivo*-like conditions.

To tackle this issue, we have successfully established and characterized the first Porcine Nasal Organoids (PNOs) cultures and used this model to study the interactions with the host and between members of the nasal microbiota. We chose *Rothia nasimurium*, a prevalent but not abundant taxon, *Moraxella pluranimalium*, a prevalent and abundant taxon, and a non-virulent strain of the pathobiont *Glaesserella parasuis,* also prevalent and abundant member of the nasal microbiota of young pigs^23,24^. In addition, we included in the study a virulent *G. parasuis* strain that can disseminate systemically and produce Glässer’s disease, a systemic inflammatory disease characterized by polyserositis^25^. With this model, we demonstrated a crosstalk between bacteria and the host, simulating the *in vivo* situation. Remarkably, we found that *R. nasimurium* modulated the secretion of interleukins and cytokines by the PNOs when co-cultured with the other bacteria. Understanding the interplay between the host with its microbiota, and the pathogens will facilitate the design of preventative and therapeutic interventions to promote respiratory health.

## RESULTS

### Generation and characterization of Porcine Nasal Organoids (PNOs)

Several lines of PNOs were established from nasal turbinates by isolation of the cells and embedding in Matrigel (basement membrane matrix), adapting published protocols for human and swine tissues ^18,20,26^. After 7-10 days in culture, PNOs were formed showing the typical cystic shape with visible lumen (Figure 1A and 1B). Moreover, beating cilia were observed inside the lumen (Supplementary Video 1) under light microscopy. PNO cultures were treated with 1% penicillin-streptomycin and 10 μg/ml of primocin to avoid bacterial growth, but they were positive for *Mycoplasma* spp. Since *M. hyorhinis* can be found in the nasal cavity of pigs, we performed a specific PCR for this pathogen and confirmed that the PNOs were positive to this species.

**Figure 1.**
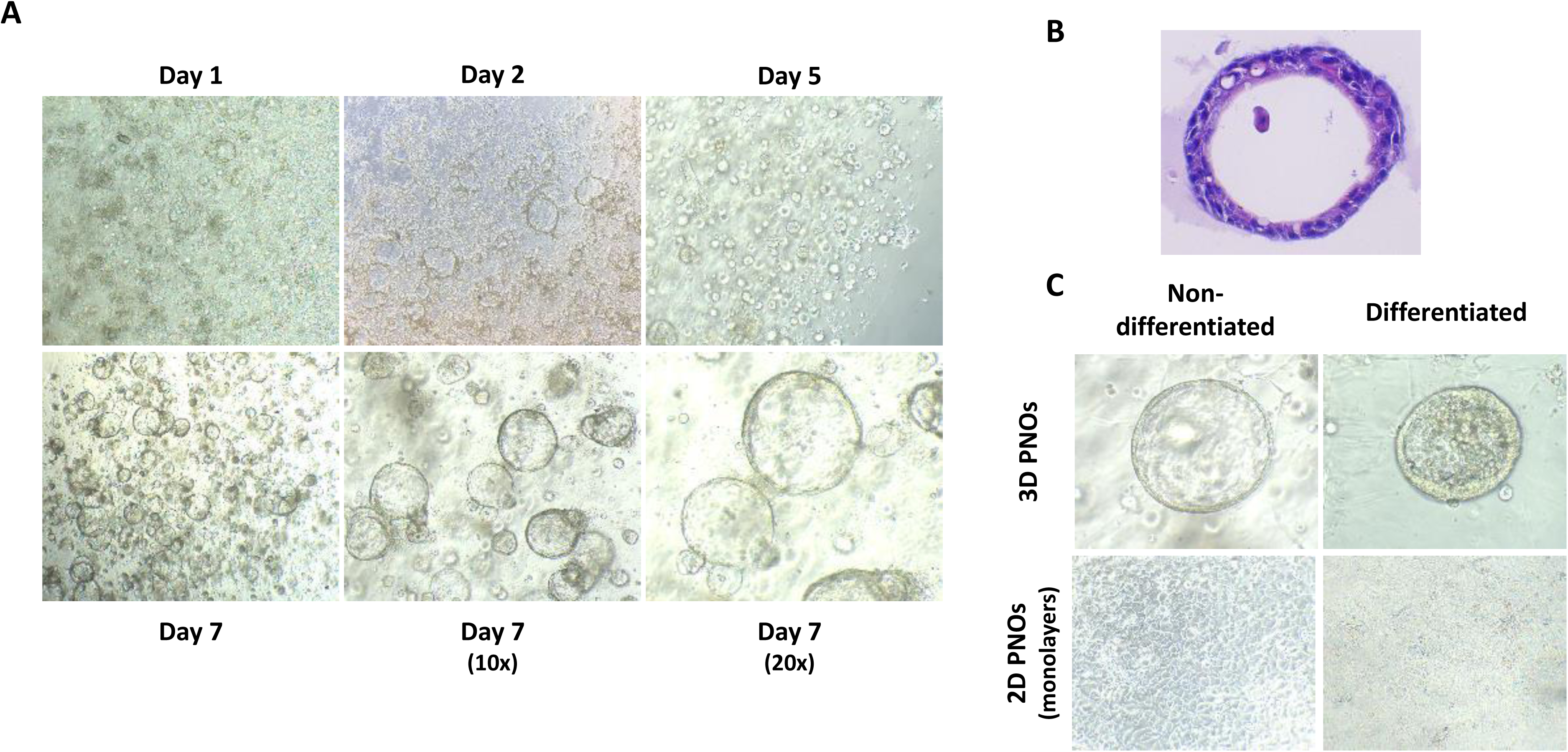
Isolation of porcine nasal organoids (PNOs) from nasal turbinates. A) Representative images of PNOs morphology at different time points by bright field microscopy at 4x, 10x and 20x magnification, respectively. B) Hematoxylin and eosin (H&E) staining of 3D paraffin-embedding PNOs. C) Comparison between differentiated and non-differentiated 3D PNOs (top) and monolayers (bottom).

To verify if the generated organoids recapitulated the histology of the nasal mucosa, organoids were expanded, differentiated and characterized. Thus, PNOs from non-differentiated 3D cultures were seeded as monolayers (2D cultures) with the apical side exposed and were cultured for 2 weeks. Monolayers were cultured in organoid growth medium (OGM) until the seventh day when the medium was switched to organoid differentiation medium (ODM) and maintained for 7 additional days (Figure 1C). PNOs were also established from nasal samples taken with cytology brushes yielding comparable PNOs. The only difference noted was that they required more time to form 3D PNO structures, specifically 2 to 3 weeks after initial adaptation to the growth medium. Early passages from non-differentiated 3D PNOs were frozen to keep a stock of the different PNO cultures and maintained viability after at least 3 freeze-thaw cycles.

To evaluate whether 3D and/or 2D PNO cultures maintained and mimicked the characteristics and the organization of the nasal respiratory epithelium, as well as the response to differentiative stimuli, the expression of several cell markers was analysed at different time points by RT-PCR and immunofluorescent assays (IFA). Similar to nasal turbinates from pigs, 3D PNOs and 2D non-differentiated and 2D differentiated PNOs expressed the markers for club cells (*scgb1a1*), goblet cells (*muc5ac*), basal cells (*tp63*), ciliated cells (*foxj1*) and tight junctions (*zo-1*). An increase in the expression of *muc5ac, zo-1* and *foxj1* genes was observed in organoids that underwent a longer period of differentiation (Figure 2A).

**Figure 2.**
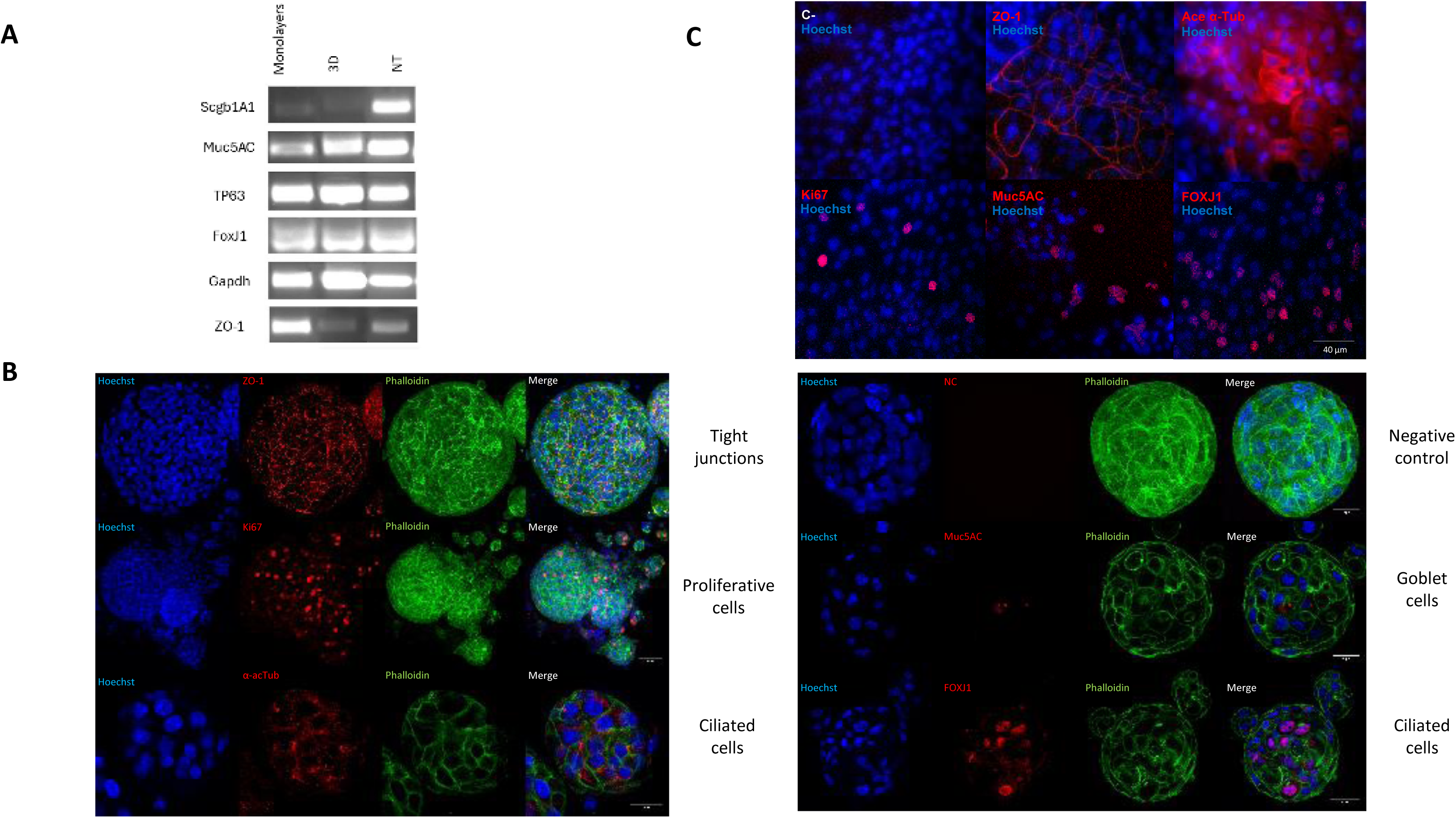
Analysis of different cell markers expressed on the PNO surface by immunofluorescent assay (IFA) and conventional PCR. Both 3D and 2D PNOs were cultured on growth medium for 7 days and then switched to differentiation medium for 7 days more and fixed with paraformaldehyde (PFA). A) mRNA expression of nasal epithelial marker genes. B) Cell marker expression in PFA-fixed differentiated 3D PNOs and C) PFA-fixed differentiated monolayers by IFA. ZO-1 protein (tight junctions) highlighted the tightness of the epithelial monolayer. Few proliferative Ki67 positive cells were detected indicating a mature epithelium. Muc5AC (goblet cells) and airway-specific markers acetylated α-tubulin and FOXJ1 (ciliated cells) were expressed at high levels. All markers were detected in red. Nuclei were counterstained with Hoechst 33342 (blue). In the 3D PNO IFA phalloidin (green) was also added to the incubation with the secondary antibody.

Gene expression of the cellular markers was confirmed by immunostaining, showing that both 3D and 2D PNO cultures did cover all major airway epithelial cell types, including proliferative cells (Ki67+), goblet cells (MUC5AC+) and ciliated cells (ACETaTUB+, FOXJ1+), and exhibited an *in-vivo* like stratification, as illustrated by the increased expression and organized localization of ZO-1 protein (Figure 2B and 2C). Few Ki67 positive cells were detected while acetylated α-tubulin appeared highly expressed in the differentiated cultures. Goblet cells were also detected. Taken together, these results suggest that the generated PNO cultures recapitulate the nasal pseudostratified airway epithelium, retaining *in vivo* morphological and functional characteristics (see Figure S1).

### Nasal microbiota members were able to colonize PNO monolayers maintaining monolayer integrity

The colonization of the host mucosal surfaces by the microbiota is fundamental in host homeostasis. To evaluate the nasal colonization with the organoid model, we selected some bacterial isolates commonly found in the nasal cavity of pigs. We used two prevalent and abundant members of the nasal microbiota: *Moraxella pluranimalium* (Mp) and *Glaesserella parasuis* (G). From the latter, we included two strains with different virulence capacities (non-virulent, nvG; and virulent, vG). Finally, *Rothia nasimurium* (Rn) was also inoculated to PNOs as a prevalent but not abundant member of the nasal microbiota. The colonization capacity was assessed through the evaluation of bacteria attachment to the organoids after overnight incubation. As a control, we also incubated the monolayers with all these strains for 2 h to analyse their adhesion capacity.

All individual strains were able to adhere and colonize the nasal organoids and were also recovered from the culture supernatants after overnight incubation (see Figure S2). *R. nasimurium* adhered (2h) to both non-differentiated and differentiated PNOs (Figure 3A and 3B). This association occurred during the first hours of incubation and did not increase in time, while Mp and both nvG and vG strains colonized the PNOs in a time dependent manner in both non-differentiated and differentiated PNOs (Figure 3A and 3B). For nvG and vG strains, adhesion at 2h was higher on differentiated than non-differentiated PNOs. Although no differences in adhesion were detected among the strains, overnight colonization by Rn was less efficient than the colonization by the other strains (Figure 3), probably related to the lower capacity of this strain to grow in the conditions of the assay, as reflected by the lower number of bacteria recovered from the supernatant (see Figure S2).

**Figure 3.**
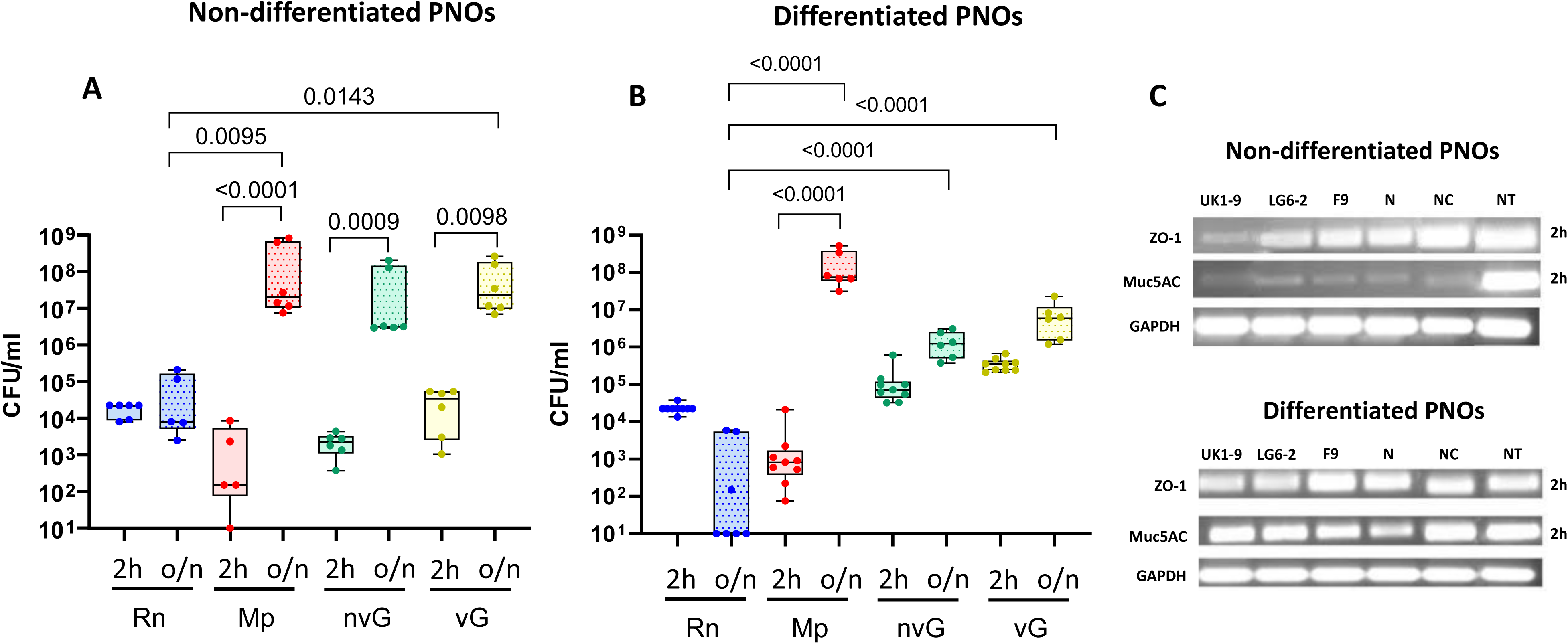
Quantity of bacteria associated with Porcine Nasal Organoids (PNOs) after 2 and overnight incubation. PNOs coming from two animals were incubated with approximately 10^5^ CFU/ml of *R. nasimurium* (Rn, in blue), 10^3^ CFU/ml of *M. pluranimalium* (Mp, in red), 10^3^ CFU/ml of non-virulent *G. parasuis* (nvG, in green) or 10^6^ CFU/ml of virulent *G. parasuis* (vG, in yellow) for 2 h (plain pattern) or overnight (o/n; dotted pattern). After incubation, non-attached bacteria were eliminated by washing, and attached bacteria were quantified by dilutions and plating after lysis of the cells with saponin. Three independent experiments were performed with duplicate wells and each dot in the plot represents individual well results. A) Non-differentiated PNOs, B) Differentiated PNOs. C) Presence of mucus (Muc5A gene, secreted by goblet cells) and the tight junctions (ZO1 gene) after 2 h of infection in non-differentiated and differentiated PNOs, respectively. Significant differences are shown in the graph as *P* values using Kruskal-Wallis multiple comparison with Benjamini, Krieger and Yekutieli post-hoc test.

### PNOs co-culturing experiments enabled to evaluate nasal microbial interactions within the microbiome and with a respiratory pathogen

*Rothia* has recently received attention for its role in host health and interaction with other members of the microbiota^27^. To explore if the PNO model can be useful for elucidating interactions between nasal microbiota members, differentiated PNOs were inoculated with combinations of the commensal *R. nasimurium* and the other bacteria under study. The colonization of Mp, nvG and vG was not affected by the presence of Rn in the co-culture (see Figure S3) and likewise, Rn colonization was not affected by the presence of any of the other strains (Figure 4A). Adhesion at 2h of either Mp or nvG in co-culture with Rn was not affected (Figure 5). However, a small reduction in the adhesion of vG was observed when co-cultured with Rn (Figure 5), which was accompanied by a reduction in the recovery of Rn from the supernatant (*P* = 0.0151; Figure 4B), and this could not be explained by competition of the strains in the culture media alone (see Figure S3). Rn 2 h adhesion was not affected by any of the strains (see Figure S4).

**Figure 4.**
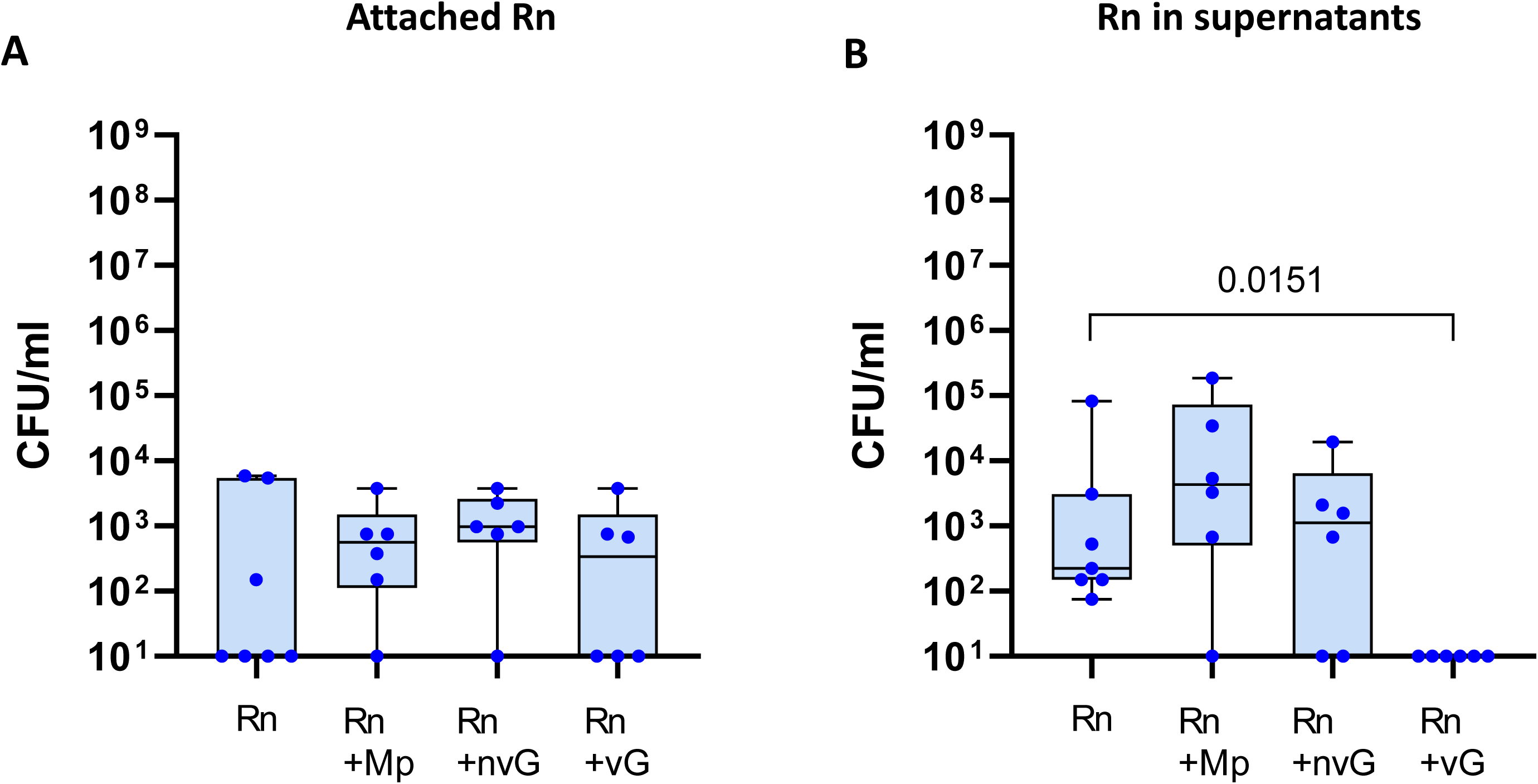
*Rothia nasimurium* load after overnight incubation with differentiated porcine nasal organoids (PNOs) individually or in co-culture with selected strains. Differentiated PNOs were inoculated with 10^5^ CFU/ml of *R. nasimurium* (Rn) alone or in combination with approximately 10^3^ CFU/ml of *Moraxella pluranimalium* (Mp), 10^3^ CFU/ml of non-virulent *Glaesserella parasuis* (nvG) or with 10^6^ CFU/ml of virulent G. parasuis (vG). After overnight incubation, attached Rn (A) and Rn in the supernatant (B) were quantified by dilutions and plating. Three independent experiments were performed with duplicate wells and each dot in the plot represents individual well results. Significant differences are shown in the graph as P values. Kruskal-Wallis multiple comparison with Benjamini, Krieger and Yekutieli post-hoc test was used to compare the bacterial concentrations.

**Figure 5.**
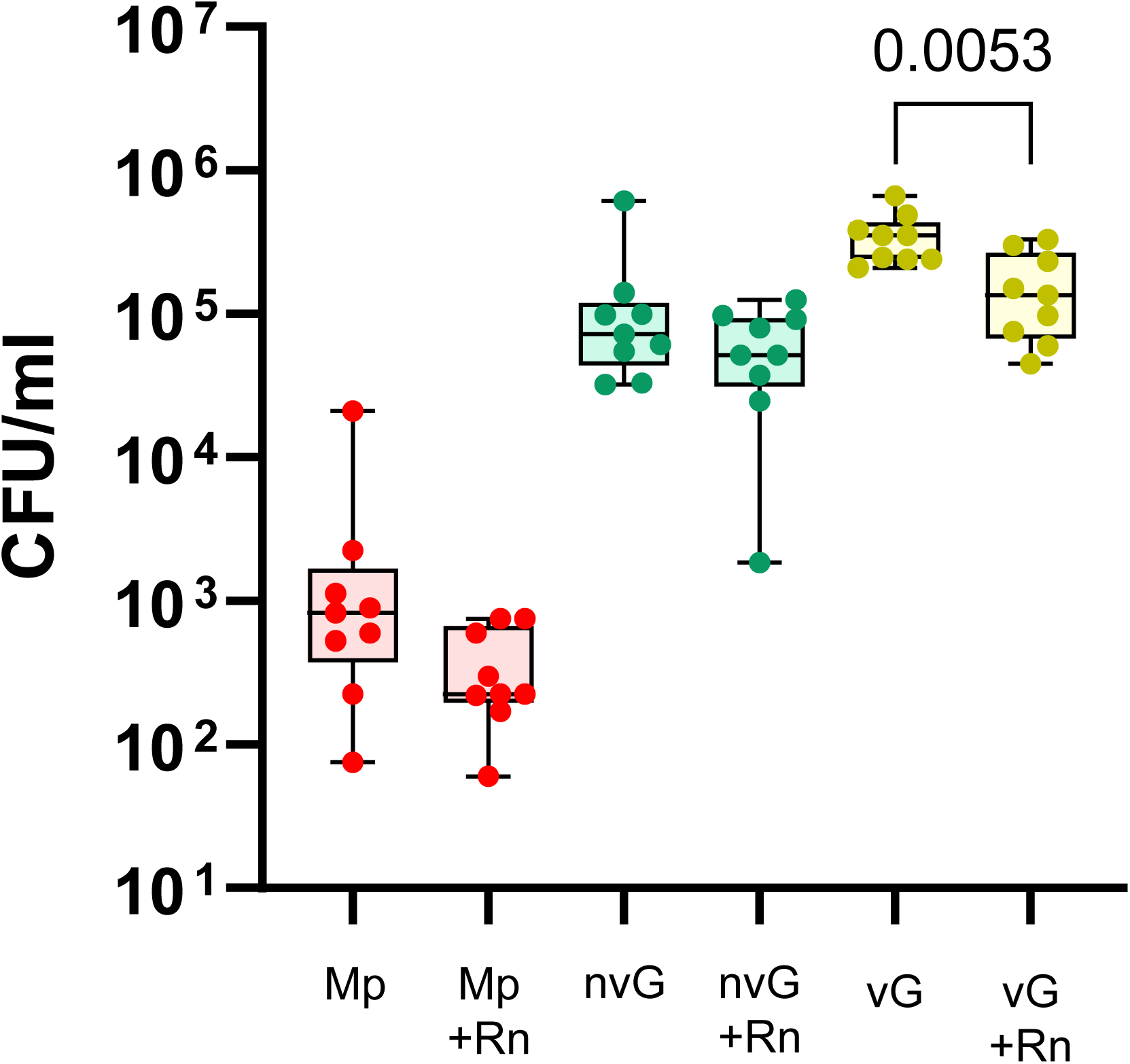
Adhesion of *Moraxella pluranimalium* and non-virulent and virulent strains of *Glaesserella parasuis* co-cultured with *Rothia nasimurium* in the Porcine Nasal Organoids (PNOs). Differentiated PNOs were incubated for 2h with approximately 10^3^ CFU/ml of *Moraxella pluranimalium* (Mp), 10^3^ CFU/ml of non-virulent *Glaesserella parasuis* (nvG) or with 10^6^ CFU/ml of virulent *G. parasuis* (vG) alone or in combination with 10^5^ CFU/ml of *R. nasimurium* (Rn). Adherent bacteria were quantified (CFU/ml) after eliminating non-attached ones by washing. Quantification of adherent Mp, nvG and vG after individual incubation or in co-culture with *Rothia nasimurium* (Rn) are shown in the graph (Mp+Rn, nvG+Rn and vG+Rn). Each dot represents a replicate from three different experiments of PNOs coming from two animals. Two replicates were included in each experiment. Significant differences are shown in the graph as P values using Welch’s t test.

Minor differences in colonization patterns were observed in non-differentiated PNOs. A reduction of Rn colonization was observed when co-cultured with both nvG and vG strains (P = 0.0410 and P = 0.0009 respectively, see Figure S5), but the adhesion at 2h was not affected (see Figure S6). In the supernatants from the co-cultures, a reduction in the number of Rn was observed when co-cultured with the vG strain (P = 0.0055), but not with the nvG, similarly to what was observed with differentiated PNOs (see Figure S7). Considering that the initial Rn inoculum was 10^5^ CFU/ml, Rn did not grow in the conditions used for PNO culture. In fact, when Rn was grown as control in fresh OGM and ODM without PNOs, no colonies were recovered after overnight incubation (see Figure S8A). However, when co-culturing Rn with Mp or both nvG and vG strains, the quantity of Rn recovered after incubation was similar to the inoculum used (see Figure S8A). By contrast, no differences between individual growth and growth with Rn in co-culture were observed in Mp when inoculated in fresh OGM and ODM (see Figure S8B), as this bacterium can grow in the same way in all conditions tested. Interestingly, when inoculating ∼10^3^ CFU/ml of nvG, no growth was detected in single culture in fresh OGM or ODM but reached to approximately 10^4^ and 10^5^ CFU/ml when co-cultured o/n with Rn in OGM and ODM respectively, indicating a metabolic cooperation between these strains (see Figure S8C). Similar results were observed with vG, as it reached a higher concentration of bacteria when co-cultured with Rn in the absence of PNOs (see Figure S8D). Overall, these results indicate that the PNOs are providing a substrate for Rn’s growth, but also that vG outcompetes Rn in these conditions. By contrast, it seemed to be a cooperation between nvG and Rn. Similar to differentiated PNOs, colonization of non-differentiated PNOs by Mp and both nvG and vG strains was not affected by the presence of Rn (see Figure S9). However, Rn reduced the initial adhesion of nvG (2 h of incubation) in the non-differentiated PNOs (P = 0.0189, see Figure S6).

### *R. nasimurium* modulated the PNOs secretion of interleukins and cytokines and had an anti-inflammatory role on PNOs

The results described above indicated that Rn did not have a direct impact on other microbiota members largely (i.e. by altering their abundance). To explore a possible interaction through immunomodulation, we quantified cytokine release by PNOs in presence/absence of Rn and the other members of the nasal microbiota. For that, multiple cytokines (IFNα, IL-1β, IL-4, IL-10, IL-17A, IL-8, IL-12, IL-6, TNFα and IFNγ) associated with inflammation and colonization were assessed. In general, two different patterns were detected in the stimulation by the individual strains of the differentiated PNOs. The first one produced by Rn and characterized by the induction of IFNγ (Figure 6A). The second one, the stimulation by the rest of the strains, characterized by a proinflammatory pattern with induction of IL-8, IL-12 and TNFα (Figure 6B, 6C and 6D respectively). Interestingly, the pro-inflammatory effect of Mp and both nvG and vG was abolished by co-incubation with Rn (Figure 6), despite the low numbers of Rn in the co-cultures (∼10^3^ CFU/ml) in comparison to the high abundance of nvG and vG (10^7^ CFU/ml for both strains) and Mp (10^8^ CFU/ml). On the contrary, the stimulation of IFNγ by Rn was not affected by incubation with the other two commensal strains (Mp and nvG), but it was abolished by the pathogenic one, the vG strain (Figure 6A).

**Figure 6.**
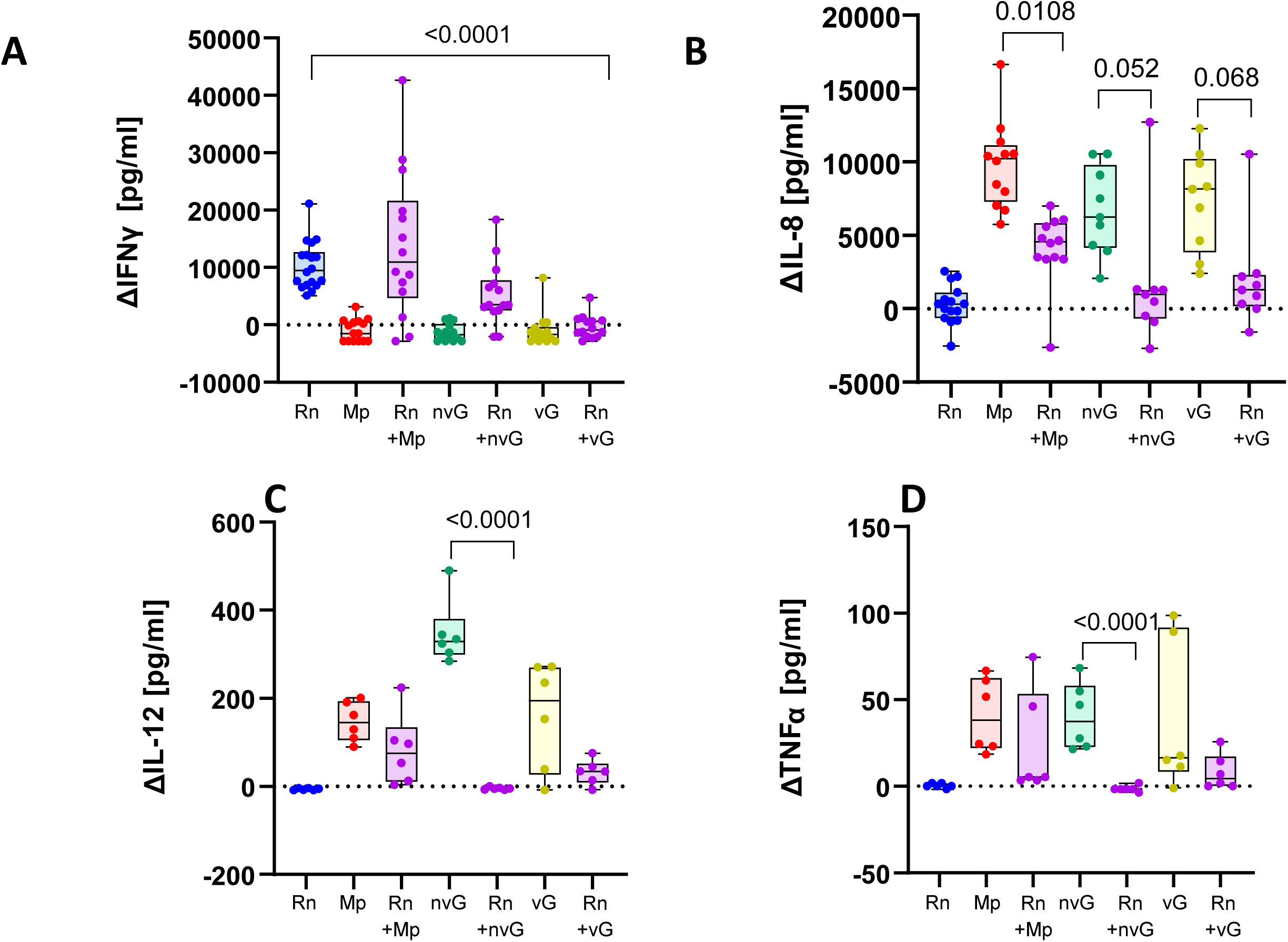
Interleukins and cytokines concentrations secreted by the differentiated Porcine Nasal Organoids (PNOs) when stimulated with the bacteria. Differentiated PNOs were incubated with 10^5^ CFU/ml of *R. nasimurium* (Rn), 10^3^ CFU/ml of *Moraxella pluranimalium* (Mp), 10^3^ CFU/ml of non-virulent *Glaesserella parasuis* (nvG) or with 10^6^ CFU/ml of virulent *G. parasuis* (vG) individually or in combinations. After overnight incubation supernatants were recovered for cytokine quantification. Results are shown as the delta of each interleukin and cytokine, subtracting the negative control (PNOs without inoculation, basal levels of the secretion) in pg/ml. A) Secretion of interleukin 8 (IL-8), B) Secretion of interleukin 12 (IL-12), C) Secretion of Tumor Necrosis Factor alpha (TNFα) and D) Secretion of Interferon gamma (IFNγ). Each dot represents a replicate from three different experiments of PNOs coming from two animals. Two replicates were included in each experiment. Kruskal-Wallis multiple comparison with Benjamini, Krieger and Yekutieli post-hoc test was used to analyze the data.

IFNα, IL-1β, IL-4, IL-10 and IL-17A (data not shown) were below the detection limits in all tested samples, suggesting that these cytokines are not released in this PNO model under the tested conditions.

Similar results in the expression of cytokines were obtained with non-differentiated PNOs (see Figure S10B and S10C respectively), with the addition of the ability of secrete IL-6 (see Figure S11).

Finally, IFNα, IL-1β, IL-4, IL-10 and IL-17A were below the detection limits in all tested samples, suggesting that t these cytokines are not released in this PNO model under the tested conditions.

## DISCUSSION

In this study, for the first time, we have established long-term cultures of porcine nasal organoids derived from nasal turbinates by isolating the basal epithelial cells. These 3D “mini-noses” exhibited a centralised lumen surrounded by a polarised airway epithelial cell layer, showing cilia oscillations with proliferative activity and tight junction function, which are physiologically similar to the pig airway epithelium *in vivo*. This model recapitulates key histological and functional aspects of the *in vivo* tissue, suggesting that it is an excellent proxy of the porcine nasal epithelium and can be used to model respiratory diseases. As a proof of concept, we used the PNOs to study host-bacterial interactions using nasal colonizers of pigs.

The culture conditions used to generate and maintain the PNOs preserved the self-renewal of nasal-resident stem cells and the stability of the organoids during the proliferation phase. This allowed us to indefinitely expand the organoid culture while recapitulating the cell diversity of the native tissue. For the differentiation stage, the composition of the medium promoted the specialisation of cells as we confirmed by PCR and IFA. Expression of FOXJ1 and MUC5AC indicated the cilia formation and mucus production, both involved in bacterial clearance. These markers, together with ZO-1 (zonula occludens-1, a tight junction protein implicated in the maintenance of cell barrier integrity) were relevant for the analysis of the host response to both colonization by nasal commensal bacteria and the infection by the respiratory pathogen. In relation to this, the presence of mucus (Muc5A gene, secreted by goblet cells) and the tight junctions (ZO1 gene) were more evident after 2 h of infection than after overnight incubation with the strains. This suggested that PNOs’ cells were more reactive to the bacteria during the first hours and reached an equilibrium after anovernight incubation. For non-differentiated PNOs this effect was not observed, suggesting that the lack of differentiation is related to less specialization. Taking this into account, we conclude that colonizers did not affect the integrity of PNO monolayers, whereas vG did disrupt the tight junctions and affected the production of mucus, damaging goblet cells and probably other cellular types, which is in agreement with other studies^28–30^.

Interactions between pathogens and members of the nasal microbiota could be essential to limit pathogen entry and their capacity to disseminate systemically^31^. For instance, in humans, several studies revealed that interactions between members of the nasal microbiota, such as *Corynebacterium spp.* or *Dolosigranulum pigrum*, interfered with *Staphylococcus aureus* colonization^32–35^. Nevertheless, the mechanisms involved in these host-microbiome interactions are difficult to study in a systematic way and could profit from a simplified and convenient model like nasal organoids. Our approach with PNOs to study bacterial colonization aligns with other works that describe the *in vivo* situation. For instance, *Rothia* is prevalent but not abundant in pig nasal swab samples, while other species as *M. pluranimalium* and *G. parasuis* showed high prevalence and abundance in this type of samples^23,24^. These findings were in apparent contradiction with the capacity of those bacterial species to grow in laboratory conditions, i.e., *R. nasimurium* grows abundantly in common bacterial media, while *M. pluranimalium* and *G. parasuis* are fastidious microorganisms that grow poorly. In our study, the initial inoculated amount was similar for all the strains, but we found that our PNO model may be mimicking the *in vivo* conditions of the microbial network and the nasal mucosa, where a high abundance of *Moraxella* and *Glaesserella* is typically detected in the nasal microbiota of pigs, together with a low abundance of *Rothia*.

Differentiated PNOs seemed to be less reactive to bacterial stimuli than the non-differentiated ones as they secreted less interleukins, especially IL-8, IL-12 and IL-6, the latter being only detected in non-differentiated PNOs. We hypothesized that this effect could be produced because differentiated PNOs have more specific cells associated to protection, as ciliated cells, and goblet cells, whose movement and secretion of mucus, respectively, could avoid intimate bacterial contact. Moreover, our results, confirm previous observations showing that Gram-negative bacteria induce pro-inflammatory cytokines IL-8, IL-12, IL-6 and TNFα, while Gram-positive tend to induce IL-12, TNFα and IFNγ^36^, supporting the value of the PNO model. IFNγ can inhibit IL-8, TNFα and IL-6 production^37–39^. In fact, IFNγ has an anti-inflammatory role when the host is infected with bacterial pathogens^37,40^, in agreement with our results and results with *Rothia mucilaginosa*. *R. mucilaginosa* was negatively correlated with pro-inflammatory markers as IL-8, IL-6 and IL-1β and demonstrated an immunomodulatory activity in a three-dimensional cell culture model of the respiratory tract in the presence of pathogens as *Pseudomonas aeruginosa*^41–43^. Similarly, Rn had an anti-inflammatory role in the PNOs by eliciting IFNγ production and interfered with the stimulation produced by the other members of the nasal microbiota, although the numbers of Rn were lower than those of the other bacteria. Overall, our results suggest that Rn might be important for the maintenance of the homeostasis with potential properties as a probiotic.

We present a novel model that recreates the typical aspects of the *in vivo* nasal epithelium based on 3D PNOs (that can be indefinitely passaged and frozen for long-term expansion) and a monolayer culture derived from them, which can be convenient for microbial network and host-microbe interaction studies. It is also important to remark the breakthrough in the generation of the pig nasal organoids using a non-invasive technique (cytological brushes) as an alternative to necropsy-derived organoids, supporting the 3R principle of animal experimentation. Additionally, we improve our understanding of the nasal microbiota colonization and the interactions between the different members, as well as the host cellular response to both bacterial colonization and infection. All the experiments presented in this study have been performed in monolayers using 96-well plates as a “*proof of concept*”. Despite monolayers recapitulated the *in vivo* structure, future improvements would include air-liquid interface (ALI) cultures to simulate apical-basal compartmentalization, and thus increase differentiation. Moreover, additional studies including more members of the nasal microbiota, as for example, a consortium, should be performed to fully understand the interaction network of the microbiota.

In conclusion, PNOs demonstrated to be a promising tool to expand the knowledge on the bacteria-bacteria and bacteria-host interactions occurring in piglets’ noses, as well as useful to study pathogen exclusion to find new alternatives to antimicrobials and reduce the use of these drugs. Besides, PNOs can be taken into consideration for studies to predict future transmissions and pandemics, as pigs are natural reservoirs for a wide variety of zoonotic pathogens.

## ACKNOWLEDGMENTS

The authors want to thank the IRTA-CReSA’s BSL2 technicians for their support in the experimental part. We also want to thank Alejandro Moreno for the *muc5ac*, *scgb1a1*, *tp63*, *foxj1* and *gapdh* primer design, and Lola Pailler for her statistical suggestions. A special thanks to Joan Repullés, head of Bioimaging Unit, for the imaging processing and analysis. This work was supported by funding from MICIU (Spanish Government) projects PID2019-106233RB-I00/AEI/10.13039/ 50110 0011033 to VA and FCF and PID2022-138657OB-I00/AEI/10.13039/ 50110 0011033 to VA and MS. LBL is supported by a FPI fellowship (PRE2020-096048/AEI/10.13039/ 50110 0011033). Finally, the authors acknowledged the support of Centres de Recerca de Catalunya (CERCA) Program.

## AUTHOR CONTRIBUTIONS

Conceptualization and design of the study: VA, MS, FCF and KK. Funding acquisition: VA, MS and FCF. Performed experiments: LBL, NCV, FT, MP and JM. Performed analyses: LBL and NCV. Supervised the analyses: VA, MS and FCF. Writing of the manuscript: LBL and NCV. Manuscript reviewing and editing: VA, MS, FCF, KK, NCV, LBL, FT, MP and JM. All authors read and approved the final manuscript.

## SUPPLEMENTAL INFORMATION

Document S1. Figures S1-S11

Video S1. Cilia oscillations in 3D PNOs.

## MATERIALS AND METHODS

### Animal experimentation and ethics approval

For the obtention of nasal turbinates, euthanasia was performed following good veterinary practices. According to European (Directive 2010/63/EU of the European Parliament and of the Council of 22 September 2010 on the protection of animals used for scientific purposes) and Spanish (Real Decreto 53/2013) normative, this procedure did not require specific approval by an Ethical Committee (Chapter I, Article 3. 1 of 2010/63/EU).

### Establishment of porcine nasal organoids (PNOs) from nasal turbinates and nasal brushes

Porcine nasal organoids were isolated from two 4-week-old piglets coming from Spanish conventional farms. Piglets were transported to IRTA-CReSA facilities where they were humanly euthanized by means of an overdose of sodium pentobarbital (Dolethal). Immediately after, the complete nasal turbinates were dissected, cut longitudinally and collected in a container with Dulbecco’s modified Eagle’s medium (DMEM/F12) complemented with HEPES, GlutaMax, Penicillin-streptomycin (P/S) and Amphotericin B (AmphB) (complete DMEM/F12). Samples were stored on ice until processing on the lab or kept at 4°C overnight for further processing of epithelial isolation on next day. After some washes with cold PBS 1x supplemented with P/S and AmphB, nasal turbinates were transferred onto a sterile plate using sterile tweezers and scalpel. The nasal mucosa tissue was peeled off and cut into small segments. Tissue samples were washed with cold PBS 1x supplemented with P/S and AmphB in a 50-ml centrifuge tube until the supernatant was clear, allowing mucosal segments to settle down to the bottom by gravity. Then, the pieces were incubated in collagenase IV (1Lmg/mL) resuspended in 10 ml of Organoid Growth Medium (OGM; Complete PneumaCult-Ex Plus Basal Medium [StemCell Technologies Cat #05040]) containing 1% of both AmphB and P/S) at 37°C for 1 h on a rocker (200 rpm). After digestion, collagenase IV was removed, and the airway debris was dissociated with 0.25% EDTA-Trypsin at 37°C for 30 min. Then, the debris was harvested by centrifugation at 300 x g at 4°C for 4 min, resuspended in complete DMEM/F12 with 10% Fetal Bovine Serum (FBS), and strained through a 100-µm strainer. Once spun at 400 x g for 5 min at 4°C, airway cell pellet was washed with 10 ml of complete DMEM/F12 and 10 ml of OGM. Finally, the medium was removed, and the cell pellet was resuspended in Matrigel (Corning 356231), seeded into a 24-well plate, and incubated for 15-30 min at 37°C with 5% CO_2_. After solidification, Matrigel droplets were cultured with 500 µl of OGM supplemented with P/S and AmphB and placed back into 37°C incubator with 5% CO_2_. The culture medium was exchanged every other day, and the nasal organoids were passaged every 7-10 days at a 1:2 or 1:3 ratio for expansion. After 2 or 3 passages, P/S and AmphB were removed from the OGM. Additionally, PNOs were tested for *Mycoplasma* by qPCR.

PNOs were also stablished from nasal brushes using cytological brushes (Covaca). The protocol followed was the same as from nasal turbinates with some modifications. Briefly, cytological brush was inserted into the nasal cavity, softly spun and then placed into a 15-ml tube containing 10 ml of cold complete DMEM/F12. Cellular material was removed using a cut-ended pipette tip and transferred to another 15-ml tube. Both tubes were centrifuged at 300 x g for 4 min at 4°C and the pellet was washed twice with cold PBS 1x supplemented with P/S and AmphB. TrypLE (Gibco) was added to this pellet, incubated for 10 min at 37°C and inactivated with 5 ml of complete DMEM/F12. After that, the cellular material was passed through a 70-µm filter to obtain the epithelial cells, spun at 300 x g for 4 min at 4°C and resuspended in Matrigel.

### 2D airway organoid monolayer cultures

Airway organoid monolayer cultures were prepared from 3D airway organoids as previously described (Ettayebi 2016). First, after culturing for 7 days, 3D airway organoids were washed by using ice-cold DMEM/F12 and dissociated with 0.05% trypsin-EDTA (Gibco) in a 37° water bath for 4 min. After incubation, trypsin was inactivated by adding DMEM/F12 containing 10% of FBS and the organoids were vigorously repeated pipetted up and down to dissociate into single cells. Then, the cell suspension was filtered through a 40-μm cell strainer (352340; Corning), centrifuged and resuspended in OGM with 10 µM rock inhibitor (Y-27632). Finally, cells were seeded at 3×10^5^ airway organoid cells in each well of a 96-well plate precoated with Matrigel. After culturing for 7 days, the 2D airway organoid monolayers were ready for the microbiota experiments. Differentiated PNOS were prepared switching the OGM to Organoid Differentiation Medium (ODM; Complete PneumaCult Airway Organoid Basal Medium [StemCell Technologies Cat #05061]) at day 7, replacing media every other day for one week more.

### Observation of cilia oscillations

The nasal mucosa organoids, both 3D and 2D models, were observed at different cultured times for cilia movement under an optical microscope (x20 and x40) and recorded by video (Supplementary Video 1).

### Fixation, paraffin-embedding, and sectioning of 3D PNOs

The protocol followed is based on a previous publication (Wiley 2017). Briefly, 3D organoids were rinsed with PBS and fixed with PFA 4% for 1h at 4°C. After several washes with PBS, organoids were centrifuged for 5 min at 1500 rpm and then were embedded in agarosa 2%, cut in small, rounded pieces and introduced into the cassettes followed by dehydration in ethanol series (80%, 95% and 100%, 15 min each at RT). The 3D organoids were then subjected to paraffin embedding and sectioning. Standard haematoxylin and eosin (H&E) staining were performed to evaluate the airway structure.

### Immunohistochemistry

Paraffin-embedded nasal turbinates from piglets were used to localize and compare the expression of the different cell types. Samples were sectioned at 2’5-µm thickness, deparaffinized and rehydrated. After dH_2_O washes, the endogenous peroxidase was blocked with methanol and 3% H_2_O_2_ for 30 min at RT. Antigen retrieval was performed at pH 6 using a sodium citrate buffer for 20 min at 98°C. Then, sections were washed and blocked for 1h in PBS-Tween 20-2% BSA (Bovine Serum Albumin) at RT. Primary antibodies ZO-1 dilution 1/300 (Invitrogen ZO1-1A12), acetylated α-tubulin dilution 1/10.000 (Proteintech 66200-1-Ig), Ki67 dilution 1/2000 (BD 550609) and 5 µg/ml FOXJ1 (Invitrogen 14-9965-80) were incubated overnight at 4°C. After three washes with PBS, Envision+System-HRP labelled Polymer anti-mouse (Dako) was added at RT for 45 min. The reaction was developed by adding the DAB substrate and sections were mounted with DPX.

### Immunofluorescence assays

PNO monolayer cultures in 96-well plates were incubated for 7 days with OGM (non-differentiated PNO monolayers) or 7 days with OGM + 7 days with ODM (differentiated PNO monolayers). Then medium was removed, monolayers washed twice with PBS 1x and fixed with PFA 4% for 1h at RT. After fixation, cells were washed twice with PBS 1x, permeabilized with 0.5% Triton-X100 for 1h at RT and blocked with BSA 3% for another hour. After that, primary antibodies were added and incubated overnight at 4°C. Next day, cells were washed 3 times with PBS and incubated with the secondary antibodies (α-mouse IgG1 antibody conjugated with ALEXA 568 or ALEXA 488 at 1/1000 dilution, mixed with phalloidin and Hoescht, for 4h at RT. After 4 h of incubation in darkness, monolayers were washed 3 times with PBS and 150 µl of PBS were finally added. The same protocol was followed for the 3D PNOs cultured 7 days with OGM (non-differentiated) or 7 days with OGM + 3 days of ODM (differentiated) but 3 washes were performed between steps for 5 min in agitation. Low-binding eppendorfs were used for the incubations. Approximately, 20 µl of 3D HIEs suspension were mounted on slides and mixed with VECTASHIELD Plus antifade mounting medium (Sigma-Aldrich). 96-well plates (monolayers) and slides (3D PNO) were observed using a fluorescence (MOTIC AE31E) and confocal microscope (Leica Stellaris 8), respectively.

### RNA extraction and RT-PCRs

PNOs were disaggregated and resuspended in DNA/RNA Shield (Zymo Research R1100-50) to preserve the RNA and frozen at -20°C until its extraction. Total RNA was extracted using the RNeasy Mini Kit (Qiagen 74104) following the manufacturer’s instructions. DNase digestion step was performed using the RNase-Free DNase Set (Qiagen 79254) as described and recommended by the manufacturer. Concentrations were measured using absorbance at 260nm (A_260_) with BioDrop DUO (BioDrop Ltdre). RNA was retrotranscribed to cDNA using a minimum concentration of 80 ng/ml of RNA using the PrimeScript™ RT Master Mix (Perfect Real Time) (Takara Bio #RR036B) kit following manufacturer’s protocol. Finally, a conventional PCR to detect club cells (Scgb1A1), goblet cells (Muc5AC), basal cells (TP63), ciliated cells (FoxJ1) and tight junctions (Zo1) was used. A housekeeping gene (gapdh) was included as a positive control of the extraction.

### Incubation of bacteria, individually and in co-culture, with non-differentiated and differentiated 2D PNOs

Porcine nasal isolates *Rothia nassimurium* UK1-9, *Moraxella pluranimalium* LG6-2 and non-virulent *Glaesserella parasuis* F9, together with virulent *G. parasuis* Nagasaki originally isolated from meningitis, were used in this study^24,44^. For the assays, the strains were grown overnight on chocolate agar at 37°C with 5% CO_2_. Next, bacterial suspensions were prepared in order to have approximately 10^3^-10^5^ CFU/ml of each strain. Inoculums were centrifuged at 16000 x *g* for 10 min and the bacterial pellet was resuspended in OGM or ODM for non-differentiated or differentiated PNO, respectively. Serial dilutions of the final inoculum were performed and plated on chocolate agar to confirm the inoculum concentration.

PNOs were prepared in 96-well plates and were inoculated with 100 µl of each individual bacterial suspension. In the case of co-cultures, 50 µl of *R. nasimurium* inoculum were mixed with 50 µl of each of the other strains in pairwise combinations (*R. nasimurium* + *M. pluranimalium*, *R. nasimurium* + non-virulent *G. parasuis* F9, and *R. nasimurium* + virulent *G. parasuis* Nagasaki) to evaluate the possible interaction between the different strains. The 96-well plates with the inoculated 2D PNOs monolayers were incubated overnight at 37°C with 5% CO_2_ to study the colonization capacity of the strains, including stimulation capacity. Afterwards, supernatants of each well were taken for bacterial count by serial dilutions and determination of cytokine secretion by PNOs (see below). Next, 2D PNOs monolayers were washed 2 times with 150 µl of PBS and were lysed with 1 % Saponin from Quillaja Bark (Sigma S7900-25G) to assess the associated bacteria. Serial dilutions of the lysate were performed and cultured on chocolate agar. All the chocolate agar plates were incubated at 37°C with 5% CO_2_ for 48 hours. Additionally, same inoculations of the bacteria to the 2D PNOs were performed but reducing the time of incubation to 2 h to study adhesion capacity. After incubation, supernatants and PNOs monolayers were processed as indicated above.

One of the infected 96-well plates was used to RNA extraction and RT-PCR analysis for the *muc5ac* and *zo-1* gene expression, presence of goblet cells and tight junction respectively, as described before.

As a growth control, bacterial inoculums were incubated with fresh OGM and ODM. Graphpad 8.3 (538) Prism software (Dotmatics, San Diego CA) was used to analyse the data. Kruskal-Wallis multiple comparison with Benjamini, Krieger and Yekutieli post-hoc test was used to compare the growth and adhesion of *R. nasimurium* co-cultured with the other strains after overnight incubation in both non-differentiated and differentiated 2D PNOs (Figure 2). Welch’s t test and Mann-Whitney test were used to compare the concentrations of non-virulent and virulent *G. parasuis* strains after 2 h of incubation in co-culture with *R. nasimurium* in both non-differentiated and differentiated PNOs (Figure 3).

### Cytokine detection in the non-differentiated and differentiated 2D PNOs

To detect cytokines, chemokines and interleukins, supernatants were taken from the 96-well plate seeded with 2D PNOs infected with the bacteria after overnight infection. Supernatants were centrifuged at 16000 x g 10 minutes to eliminate the bacterial cells and frozen at –80°C until its use. Negative controls from non-inoculated PNOs were also included.

Cytokine & Chemokine 9-Plex Porcine ProcartaPlex™ Panel 1 assay (ThermoFisher Scientific #EPX090-60829-901) was used following manufacturer’s protocol to detect Interferon alpha (IFNα), interferon gamma (IFNγ), interleukin 1 beta (IL-1β), interleukin 10 (IL-10), interleukin 12 (IL-12), interleukin 4 (IL-4), interleukin 6 (IL-6), interleukin 8 (IL-8 [CXCL8]) and Tumour Necrosis Alpha (TNFα). Standards were prepared with the growth media of the PNOs.

In parallel, interleukin 8 (IL-8) secretion was assessed with an Enzyme-Linked Immunosorbent Assay (ELISA) following the manufacturer’s protocol ((R&D system #DY535). The coat was made with 1:180 dilution of 360 μg/ml IL-8 capture antibody in PBS 1x. 300 μl of block buffer (1% Bovine Serum Albumin, BSA in PBS 1x) were added and incubated 1 h 50 rpm at RT. Samples were added using a 1:50 dilution in the reagent diluent (0,1%BSA, 0,05% Tween20 in PBS 1x). A 1:180 dilution in the reagent diluent of 5,4ug/ml detection antibody was used.

Interferon gamma (IFNγ) secretion was also assessed with an ELISA following manufacturer’s protocol (Swine IFN gamma Polyclonal Antibody, PB0157S-100; Swine IFN gamma Polyclonal Antibody Biotinylated, PBB0269S-050; Swine IFN gamma [Yeast-derived Recombinant Protein], RP0126S-005;

Kingfisher Biotech, Inc). Samples were added 1:5 diluted in the same media used for the PNOs growthIL-10 and IL-17A ELISAs were tested as described for IFNγ.

Kruskal-Wallis multiple comparison with Benjamini, Krieger and Yekutieli post-hoc test was applied to analyze the data.

