## Supplemental Information for "Porcine Nasal Organoids as a model to study the interactions between the swine nasal microbiota and the host"

FOXJ1
40x
(ciliated
cells)

Acet.
α-tubulin
40x
(cillia)

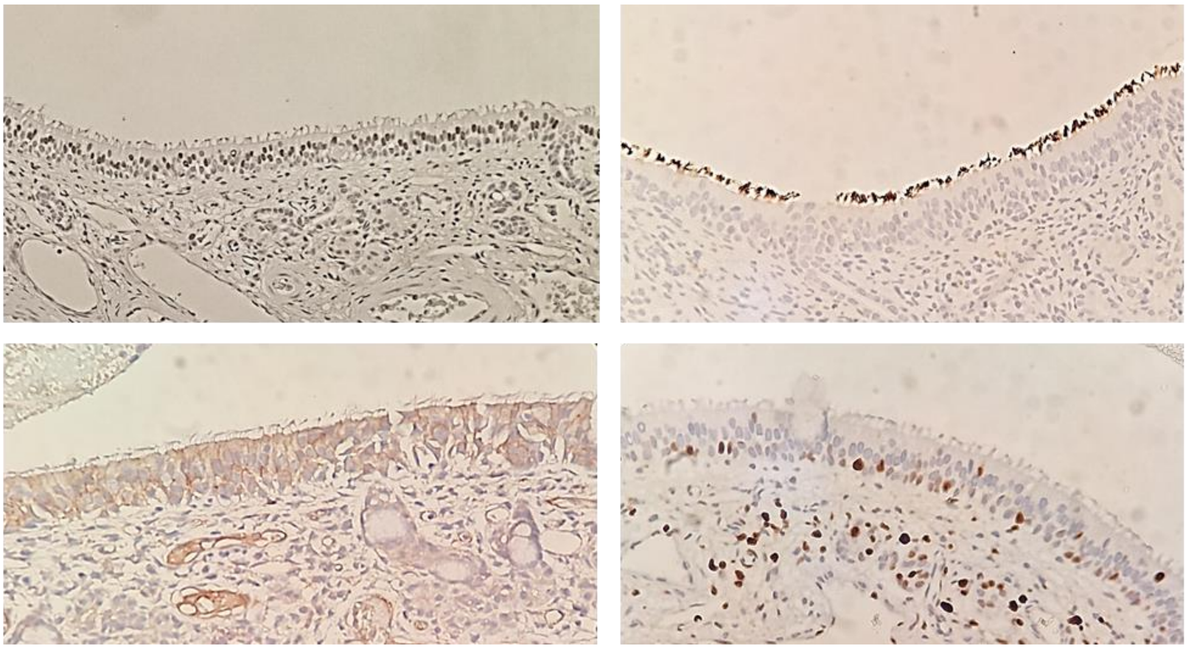

ZO-1
40x
(tight
junctions)

Ki67
40x
(cellular
proliferation)

**Figure S1. Characterization and cell marker expression in paraffin-embedded nasal turbinate from pigs by IHQ.** Nasal turbinate tissue was used for immunohistochemistry staining. Antibodies against FOXJ1 and acetylated α-tubulin (ciliated cells, left and right upper panel, respectively), ZO-1 (tight junctions, left lower panel), and Ki67 (proliferative cells) were used to visualize the different cells markers expressed on the nasal airway epithelium from pigs. Images are representative of at least five similar images.

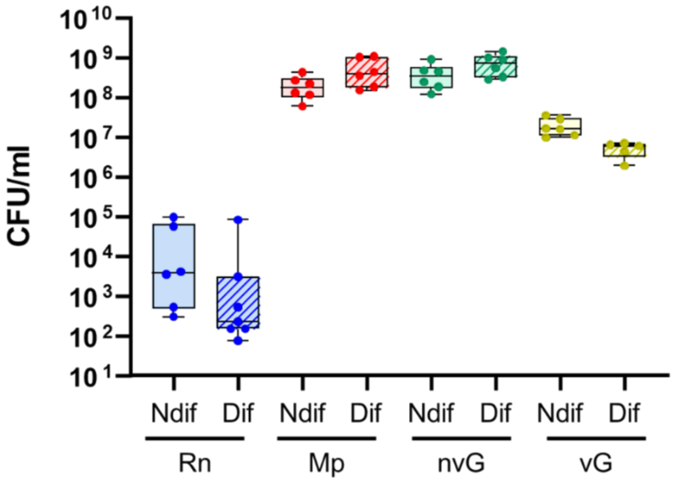


**Figure S2. Bacteria found in the supernatant of non-differentiated and differentiated Porcine Nasal Organoids (PNOs) after overnight incubation.** Non-differentiated (Ndif, plain pattern) and differentiated (Dif, stripped pattern) PNOs were overnight incubated with approximately $10^4$-$10^5$ CFU/ml of *R. nasimurium* (Rn, in blue), $10^3$ CFU/ml of *M. pluranimalium* (Mp, in red), $10^3$ CFU/ml of non-virulent *G. parasuis* (nvG, in green) or $10^5$-$10^6$ CFU/ml of virulent *G. parasuis* (vG, in yellow). After incubation, bacteria present in the supernatant were quantified by dilutions and plating. Each dot represents a replicate from three different experiments of PNOs coming from two animals. Two replicates were included in each experiment.

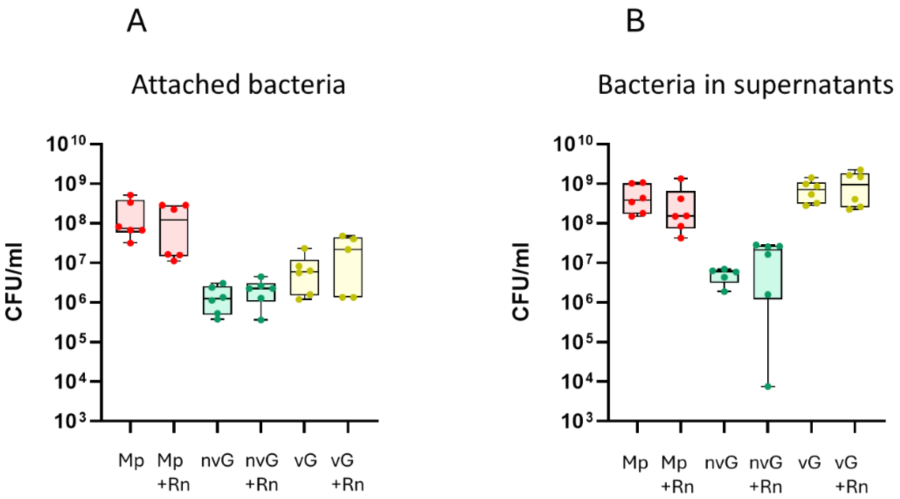


**Figure S3. Co-culture with Rothia nasimurium in the Porcine Nasal Organoids (PNOs) after overnight incubation.** Differentiated PNOs were incubated overnight with approximately $10^3$ CFU/ml of *Moraxella pluranimalium* (Mp), $10^3$ CFU/ml of non-virulent *Glaesserella parasuis* (nvG) or with $10^6$ CFU/ml of virulent *G. parasuis* (vG) individually or in combination with $10^5$ CFU/ml of *R. nasimurium* (Rn). A) As a measure of colonization capacity, attached Mp (in red), nvG (in green) and vG (in yellow) were quantified after individual incubation or co-cultured with Rn. B) Quantification of Mp (red), nvG (green) and vG (yellow) in the supernatant. Each dot represents a replicate from three different experiments of PNOs coming from two animals. Two replicates were included in each experiment.

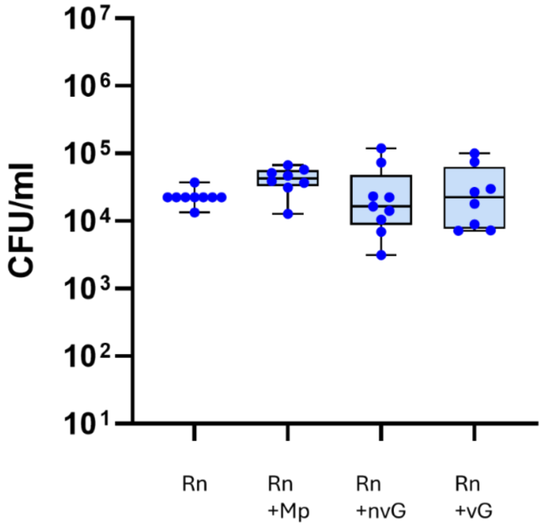


**Figure S4. *Rothia nasimurium* adhesion after 2 h of incubation with differentiated porcine nasal organoids (PNOs) individually or in co-culture with selected strains.** Differentiated PNOs were inoculated with $10^5$ CFU/ml of *R. nasimurium* (Rn) alone or in combination with approximately $10^3$ CFU/ml of *Moraxella pluranimalium* (Mp), $10^3$ CFU/ml of non-virulent *Glaesserella parasuis* (nvG) or with $10^6$ CFU/ml of virulent *G. parasuis* (vG). After 2 h of incubation, attached Rn was quantified by dilutions and plating, after washing to eliminate unbound bacteria. Three independent experiments were performed with duplicate wells and each dot in the plot represents individual well results.

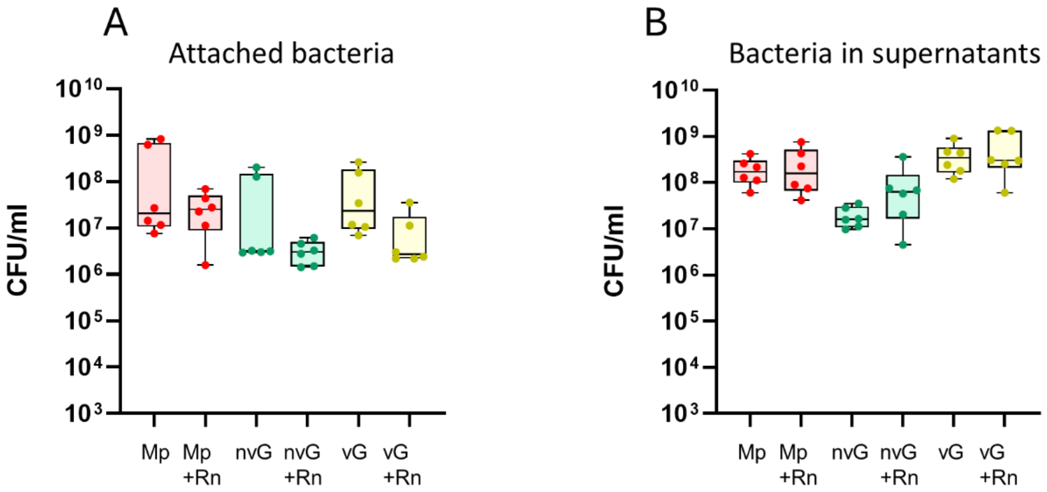


**Figure S5. Co-culture with *Rothia nasimurium* in the non-differentiated Porcine Nasal Organoids (PNOs) after overnight incubation.** Non-differentiated PNOs were incubated overnight with approximately $10^3$ CFU/ml of *Moraxella pluranimalium* (Mp), $10^3$ CFU/ml of non-virulent *Glaesserella parasuis* (nvG) or with $10^5$ CFU/ml of virulent *G. parasuis* (vG) individually or in combination with $10^4$ CFU/ml of *R. nasimurium* (Rn). A) As a measure of colonization capacity, attached Mp (in red), nvG (in green) and vG (in yellow) were quantified after individual incubation or co-cultured with Rn. B) Quantification of Mp (red), nvG (green) and vG (yellow) in the supernatant. Each dot represents a replicate from three different experiments of PNOs coming from two animals. Two replicates were included in each experiment.

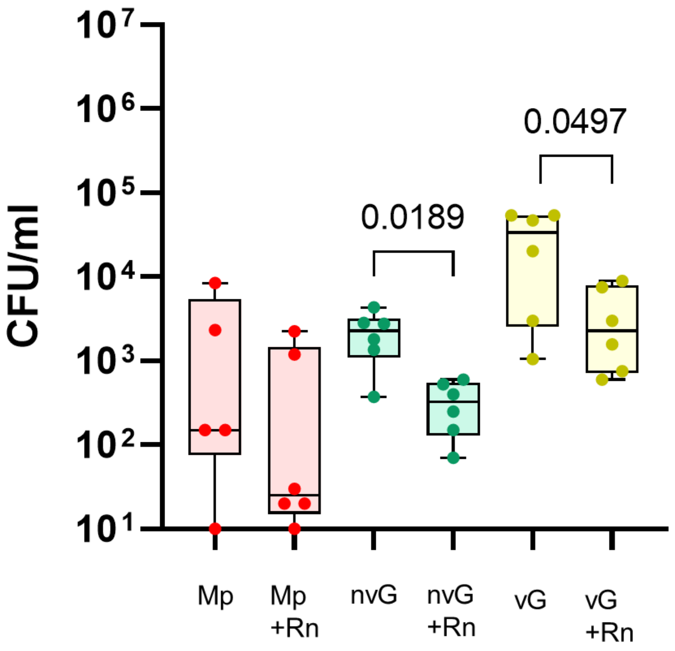


**Figure S6. Adhesion of *Moraxella pluranimalium* and non-virulent and virulent strains of *Glaesserella parasuis* co-cultured with *Rothia nasimurium* in non-differentiated Porcine Nasal Organoids (PNOs).** Non-differentiated PNOs were incubated for 2h with approximately $10^3$ CFU/ml of *Moraxella pluranimalium* (Mp), $10^3$ CFU/ml of non-virulent *Glaesserella parasuis* (nvG) or with $10^5$ CFU/ml of virulent *G. parasuis* (vG) alone or in combination with $10^5$ CFU/ml of *R. nasimurium* (Rn). Adherent bacteria were quantified (CFU/ml) after eliminating non-attached ones by washing. Quantification of adherent Mp (in red), nvG (in green) and vG (in yellow) after individual incubation or co-cultured with Rn are shown. Each dot represents a replicate from three different experiments of PNOs coming from two animals. Two replicates were included in each experiment. Significant differences are shown in the graph as P values using Welch's t test.

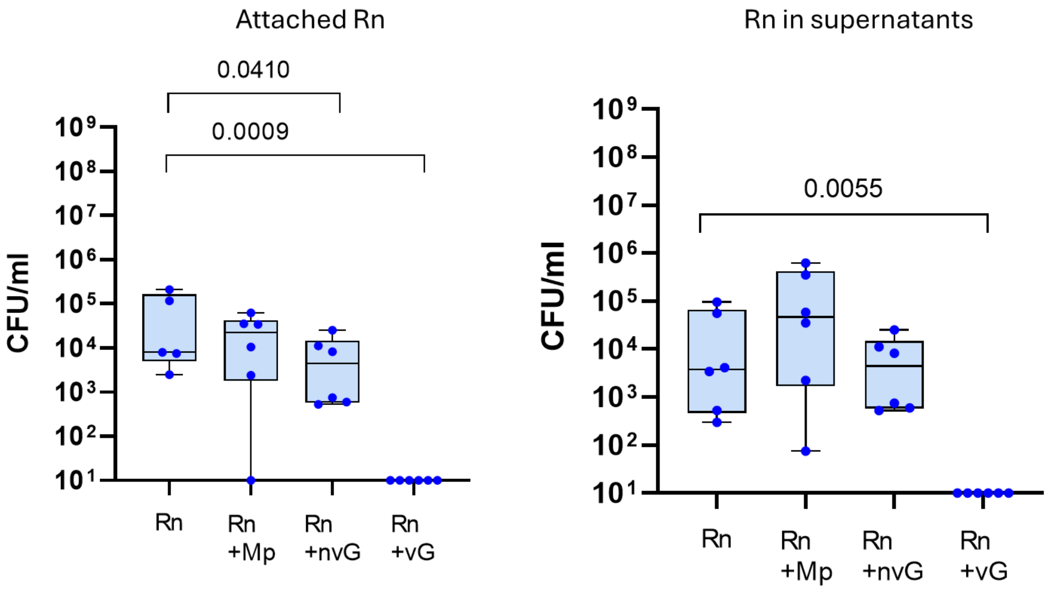


**Figure S7. *Rothia nasimurium* load after overnight incubation with non- differentiated porcine nasal organoids (PNOs) individually or in co-culture with selected strains.** Non-differentiated PNOs were inoculated with $10^4$ CFU/ml of *R. nasimurium* (Rn) alone or in combination with approximately $10^3$ CFU/ml of *Moraxella pluranimalium* (Mp), $10^3$ CFU/ml of non-virulent *Glaesserella parasuis* (nvG) or with $10^5$ CFU/ml of virulent *G. parasuis* (vG). After overnight incubation, attached Rn (A) and Rn in the supernatant (B) were quantified by dilutions and plating. Three independent experiments were performed with duplicate wells and each dot in the plot represents individual well results. Significant differences are shown in the graph as P values. Kruskal-Wallis multiple comparison with Benjamini, Krieger and Yekutieli post-hoc test was used to compare the bacterial concentrations.

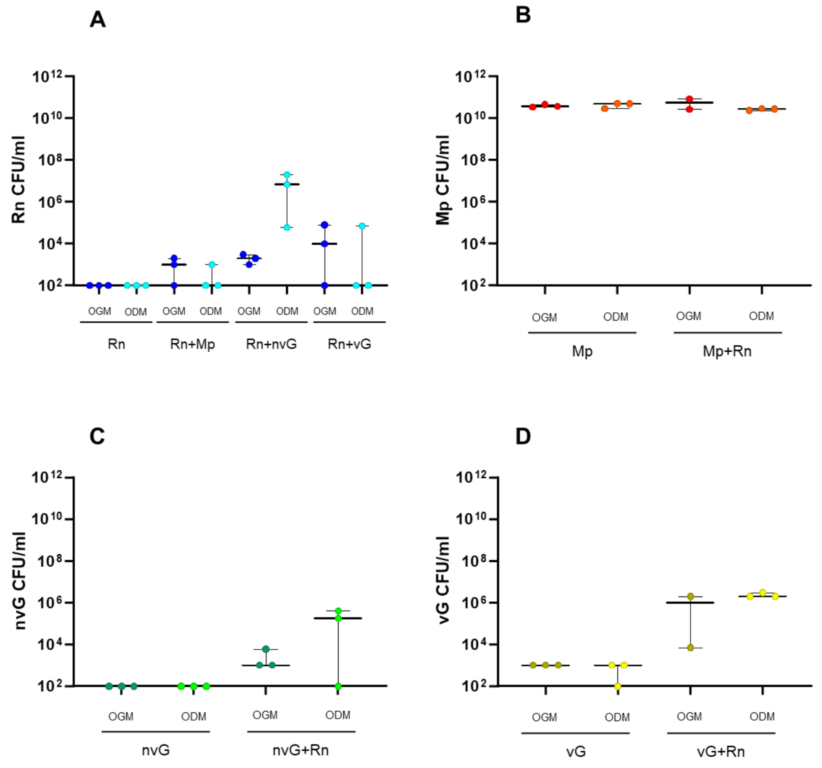


**Figure S8. Overnight co-culture of the bacterial strains in fresh organoid media.** Bacterial growth (CFU/ml) after overnight incubation in fresh Organoid Growth Media (OGM) and fresh Organoid Differentiation Media (ODM). A) *Rothia nasimurium* (Rn) individually grown as well as in co-culture with *Moraxella pluranimalium* (Mp, Rn+Mp), non-virulent *Glaesserella parasuis* strain (nvG, Rn+nvG) and virulent *G. parasuis* strain (vG, Rn+vG) in OGM (dark blue) and ODM (turquoise). B) Mp individually grown and in co-culture with Rn (Mp+Rn) in OGM (red) and ODM (orange). C) nvG individually grown and in co-culture with Rn (nvG+Rn) in OGM (dark green) and ODM (light green). D) vG individually grown and in co-culture with Rn (vG+Rn) in OGM (dark yellow), ODM (light yellow).

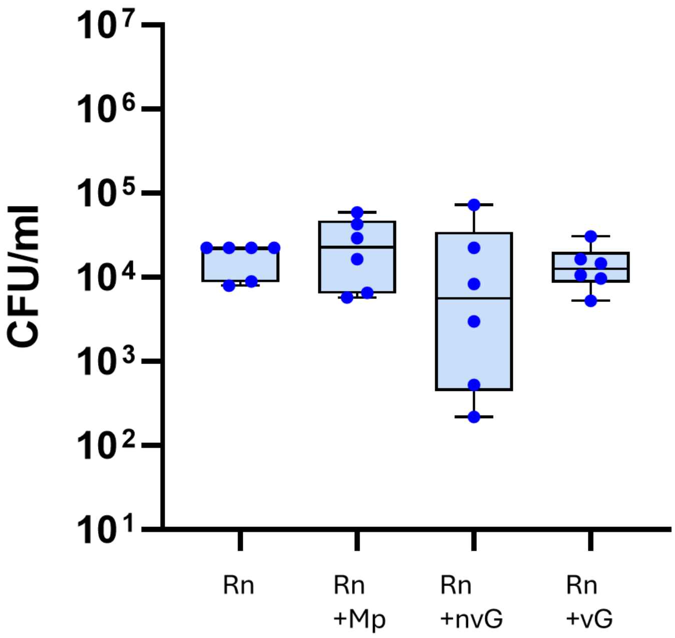


**Figure S9. *Rothia nasimurium* adhesion after 2 h of incubation with non-differentiated porcine nasal organoids (PNOs) individually or in co-culture with selected strains.** Non-differentiated PNOs were inoculated with $10^4$ CFU/ml of *R. nasimurium* (Rn) alone or in combination with approximately $10^3$ CFU/ml of *Moraxella pluranimalium* (Mp), $10^3$ CFU/ml of non-virulent *Glaesserella parasuis* (nvG) or with $10^5$ CFU/ml of virulent *G. parasuis* (vG). After 2 h of incubation, attached *Rn* was quantified by dilutions and plating after washing to eliminate unbound bacteria. Three independent experiments were performed with duplicate wells and each dot in the plot represents individual well results.

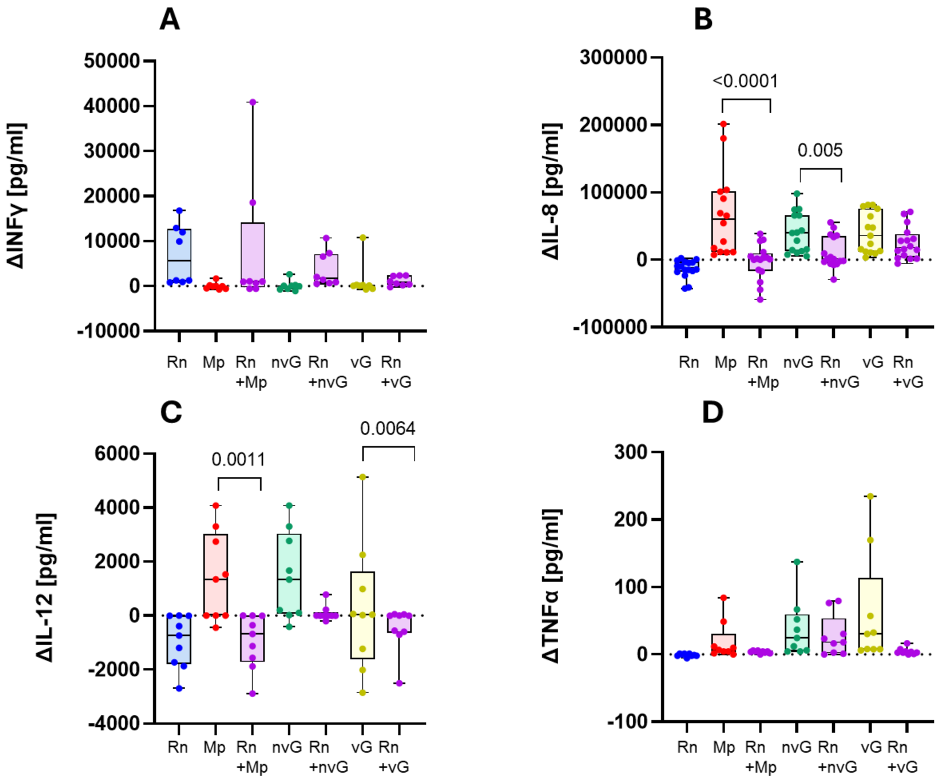


**Figure S10. Interleukins and cytokines concentrations secreted by the non-differentiated Porcine Nasal Organoids (PNOs) when stimulated with the bacteria.** Non-differentiated PNOs were incubated with $10^4$ CFU/ml of *R. nasimurium* (Rn), $10^3$ CFU/ml of *Moraxella pluranimalium* (Mp), $10^3$ CFU/ml of non-virulent *Glaesserella parasuis* (nvG) or with 105 CFU/ml of virulent *G. parasuis* (vG) individually or in combinations. After overnight incubation supernatants were recovered for cytokine quantification. Results are shown as the delta of each interleukin and cytokine, subtracting the negative control (PNOs without inoculation, basal levels of the secretion) in pg/ml. A) Secretion of interleukin 8 (IL-8), B) Secretion of interleukin 12 (IL-12), C) Secretion of Tumor Necrosis Factor alpha (TNFα) and D) Secretion of Interferon gamma (INFγ). Each dot represents a replicate from three different experiments of PNOs coming from two animals. Two replicates were included in each experiment. Kruskal-Wallis multiple comparison with Benjamini, Krieger and Yekutieli post-hoc test was used to analyze the data.

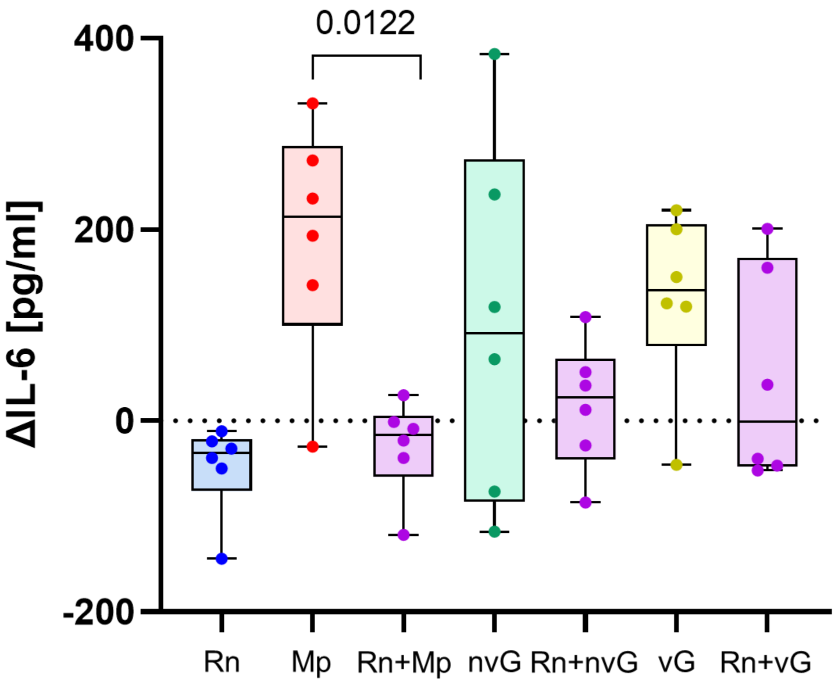


**Figure S11. Interleukin 6 (IL-6) concentrations by the non-differentiated Porcine Nasal Organoids (PNOs) when stimulated with the bacteria.** Non-differentiated PNOs were incubated with $10^4$ CFU/ml of *Rothia nasimurium* (Rn), $10^3$ CFU/ml of *Moraxella pluranimalium* (Mp), $10^3$ CFU/ml of non-virulent *Glaesserella parasuis* (nvG) or with $10^5$ CFU/ml of virulent *G. parasuis* (vG) individually or in combinations. After overnight incubation supernatants were recovered for cytokine quantification. Results are shown as the delta of interleukin 6 (IL-6), subtracting the negative control (PNOs without inoculation, basal levels of the secretion) in pg/ml. Each dot represents a replicate from three different experiments of PNOs coming from two animals. Two replicates were included in each experiment. Kruskal-Wallis multiple comparison with Benjamini, Krieger and Yekutieli post-hoc test was used to analyze the data.
